# Enhancing Collaborative Neuroimaging Research: Introducing COINSTAC Vaults for Federated Analysis and Reproducibility

**DOI:** 10.1101/2023.05.08.539852

**Authors:** Dylan Martin, Sunitha Basodi, Sandeep Panta, Kelly Rootes-Murdy, Paul Prae, Anand D. Sarwate, Ross Kelly, Javier Romero, Bradley T. Baker, Harshvardhan Gazula, Jeremy Bockholt, Jessica Turner, Nathalia B. Esper, Alexandre R. Franco, Sergey Plis, Vince D. Calhoun

## Abstract

Collaborative neuroimaging research is often hindered by technological, policy, administrative, and methodological barriers, despite the abundance of available data. COINSTAC is a platform that successfully tackles these challenges through federated analysis, allowing researchers to analyze datasets without publicly sharing their data. This paper presents a significant enhancement to the COINSTAC platform: COINSTAC Vaults (CVs). CVs are designed to further reduce barriers by hosting standardized, persistent, and highly-available datasets, while seamlessly integrating with COINSTAC’s federated analysis capabilities. CVs offer a user-friendly interface for self-service analysis, streamlining collaboration and eliminating the need for manual coordination with data owners. Importantly, CVs can also be used in conjunction with open data as well, by simply creating a CV hosting the open data one would like to include in the analysis, thus filling an important gap in the data sharing ecosystem. We demonstrate the impact of CVs through several functional and structural neuroimaging studies utilizing federated analysis showcasing their potential to improve the reproducibility of research and increase sample sizes in neuroimaging studies.

## 1 INTRODUCTION

In recent years, neuroimaging has seen a growing emphasis on data sharing and collaborative research, as evidenced by the development of new standards (e.g., Brain Imaging Data Structure (BIDS) Gorgolewski et al. (2016)), open-source software tools, and data repositories. Neuroinformatics consortia such as Enhancing NeuroImaging Genetics through Meta-Analysis consortium (ENIGMA) Thompson et al. (2014), and data repositories such as OpenNeuro Markiewicz et al. (2021) and National Institutes of Health Data Archive ^1^, were created to facilitate analysis of data and combining data from multiple sites. Pooling data from many studies allows for larger sample sizes that produce more statistical power Andrade (2020); Biswal et al. (2010). Though the quantity of neuroimaging data is increasing, there are still barriers to collaboration in the form of technological, policy, administrative, and methodological constraints that can negatively affect data accessibility.

In this section, we discuss in detail some of the challenges associated with collaborative analysis, particularly in centralized approaches, where the data need to be pooled in one location to perform an analysis. We also discuss COINSTAC, a tool built on the principles of federated analysis to enable analysis without the need to centralize data.

### 1.1 Technological Challenges

Technological constraints, such as storage space, download speed, and processing power, play a significant role in the feasibility of performing collaborative analyses on large datasets Homer et al. (2008); McGuire et al. (2011) such as neuroimaging data. Existing data repositories can contain high-resolution neuroimaging files covering tens of thousands of subjects, with sizes ranging from megabytes to multiple petabytes. Downloading the MPI-Leipzig Mind-Brain-Body dataset Babayan et al. (2020-07-22) (369.78GB) at the global median download speed of 76.32Mbps^2^ onto a modern MacBook Pro with 512GB of storage^3^ would take 11 hours and 33 minutes, consuming 72.2 percent of the machine’s total storage space. The requirements for storage space and download time can increase when an analysis involves aggregating multiple large datasets. Additionally, processing power may be a limiting factor for performing computations, particularly when certain types of analyses are designed to run on specific hardware like GPUs, which can demand resources beyond the capacity of smaller research groups or institutions with limited budgets.

### 1.2 Policy and Privacy Challenges

Due to the potentially sensitive nature of neuroimaging datasets, their use in collaborative analysis is often restricted by policies intended to preserve privacy. Collaboration methods include aggregating data in a centralized repository or using Data Usage Agreements (DUAs) Thompson et al. (2014) Thompson et al. (2017). These methods can be cumbersome and, in some cases, insufficient. DUAs may take months or even years to approve without any guarantee of the data’s utility. Data sharing might be limited by law, policy, or proprietary restrictions, largely driven by re-identification concerns. In situations where only summary data can be shared, differences in analysis methodology may result in inconsistent measures for meta-analysis Rootes-Murdy et al. (2022).

### 1.3 Administrative Challenges

Administrative challenges can arise when collaborating on an analysis, as various steps demand researchers’ time and attention. These steps may include communicating between agencies, formulating and signing data-sharing agreements, agreeing on data preparation and analysis processes, procuring technical resources, monitoring and auditing processes, performing data transfer, initiating computations, disseminating results of analyses and so on.

The efficiency of collaborative analysis is influenced by how quickly these manual steps are executed. Synchronized availability of researchers can present a barrier to the collaboration process. When researchers work asynchronously, each step in a serial process requiring manual interaction introduces potential delays. This can be particularly challenging when researchers are distributed across multiple time zones or have limited time to perform manual tasks. Furthermore, researchers’ availability may be constrained by the need for expertise and authority, such as having the authority to sign a data-sharing agreement or the technical expertise to run the appropriate Python script against a dataset. Often, these manual steps must be executed for each new analysis, which can slow down and even impede collaborative analysis. By addressing these administrative barriers, research teams can more effectively collaborate and streamline their analysis processes, ultimately contributing to the advancement of neuroimaging research.

### 1.4 Methodological differences

Variability in methodological approaches to data processing and analysis can make reproducing studies challenging Vogt (2023). To validate results, researchers must adhere to the exact methodology used in the original study, which necessitates clear communication of the specific methods employed. However, as methods are often chosen on a case-by-case basis, replicating studies can be time-consuming and difficult Esteban et al. (2019), and sometimes even impossible. Moreover, when multiple studies adopt different methodologies, combining their results meaningfully becomes challenging, hindering the execution of meta-analyses.

To overcome these barriers, we introduce COINSTAC ^4^, a tool that supports federated analysis for neuroimaging data.

### 1.5 Federated analysis using COINSTAC

Federated analysis (also federated learning, or decentralized analysis) Rootes-Murdy et al. (2022); Kairouz et al. (2021); Plis et al. (2016) allows for multiple datasets to be used in analyses without source data being directly shared. Instead, data holders run computations on their local data and only share the outputs, which are often group-level data derivatives or summary statistics. For example, sites may compute an average or other summary on their local data and send that information. Typically, these summaries are much smaller, meaning that not only are the source data not shared, and the technical challenges associated with the transfer of datasets are removed. A second potential benefit is additional privacy guarantees for the data holders. From a purely policy perspective, datasets are analyzed without being moved from their original location and data holders can determine which computations are and are not allowed on their data. From a technical perspective, strong end-to-end encryption can prevent third parties from acquiring the data derivatives. Depending on the trust model, additional privacy protections are possible, including emerging technologies like secure multiparty computation and differential privacy Dwork and Roth (2013); Heikkilä et al. (2020); Imtiaz et al. (2021); Bonawitz et al. (2017, 2016); Senanayake et al. (2022)

The Collaborative Informatics and Neuroimaging Suite Toolkit for Anonymous Computation (COINSTAC)^5^ Plis et al. (2016); Ming et al. (2017); Gazula et al. (2020, 2023); Turner et al. (2022) is a tool developed to support federated analysis specifically for neuroimaging data by overcoming the aforementioned barriers to collaboration through the use of federated analysis and standardization of collaboration methods. COINSTAC enables researchers to run decentralized neuroimaging analyses to perform larger collaborative studies Rootes-Murdy et al. (2022); Turner et al. (2022).

The COINSTAC desktop application provides an easy-to-use graphical user interface (GUI) for coordinating and executing federated analysis pipelines among multiple collaborators. Image preprocessing and a variety of univariate and multivariate approaches (e.g., VBM regression, group ICA) can be completed within the app.

One limitation of the original implementation of COINSTAC is that it requires synchronized coordination Jwa and Poldrack (2022), users have to coordinate among data owners to confirm their systems are online, that the data are organized within the same structure and that the data are mapped properly within the COINSTAC system. The need for a centralized coordinator can delay contingent analyses.

In this paper, we address this limitation by showcasing a method for hosting both private and public datasets where the datasets are persistently accessible for analysis using COINSTAC without the need for synchronized effort from data owners. Analysis of public datasets is made more accessible by removing the need to find, download, preprocess, and prepare datasets for analysis. We provide curated data vaults for various openly available neuroimaging data which COINSTAC users can simply include in their analyses. Private dataset access can be restricted to a pre-approved list of computations. Standardizing access to data vaults in the COINSTAC system simplifies analysis, optimizes computational performance, and promotes the reusability of neuroimaging datasets.

## 2 METHOD

In this section, we discuss COINSTAC and the extension of the COINSTAC framework with the addition of vaults, their architecture, and various use-cases they enable. All code for COINSTAC and COINSTAC Vaults can be found in the COINSTAC Github repository^6^.

### 2.1 COINSTAC

To understand how Vaults improve the workflow of federated analysis in COINSTAC, we will describe the COINSTAC system and how it is used.

The main components of the COINSTAC system are: the desktop application, the central server, and computation containers. The desktop application provides a graphical user interface (GUI) and manages local computation containers used to participate in federated analyses. The central server manages the central database and runs the containers that act as the inner node in federated analyses.

In the COINSTAC desktop application, users join collections of users called “consortia” to collaborate on an analysis pipeline. A consortium is a group formed by individual COINSTAC users, each with their machine that is capable of being a node in a federated analysis pipeline. Each member within a consortium will act as a node in the federated analysis group by running local computations inside of a container on their system.

The following is how a researcher would use the COINSTAC user interface to create a consortium and run a federated analysis pipeline:

- Log in as a user
- Join (as a member) or create (as an owner) a consortium
- Configure a set of computations (a pipeline) to be performed by a consortium
- Map their local data to the pipeline
- Initiate the pipeline (a run)
- View the results of the pipeline run

### 2.2 COINSTAC Vaults

#### 2.2.1 Purpose and high level overview

The Vaults system is an extension of the COINSTAC platform that allows datasets to be persistently available for participation in federated analyses without requiring manual action from data owners apart from the initial setup. COINSTAC consortium owners can independently add Vaults members to their consortia, allowing vault datasets to participate in federated analyses without the need for coordination between consortia owners and Vault data owners. The Vault client allows datasets to be made available to the larger COINSTAC ecosystem, giving the ability for others to run pipelines using the Vault’s data without it ever leaving its respective system.

#### 2.2.2 Using the GUI to add a Vault to a consortium and run an analysis

Vault clients can be added to a consortium by a consortium owner without any action required from the owner of the Vault data, as shown in Fig. 1.

**Figure 1.**
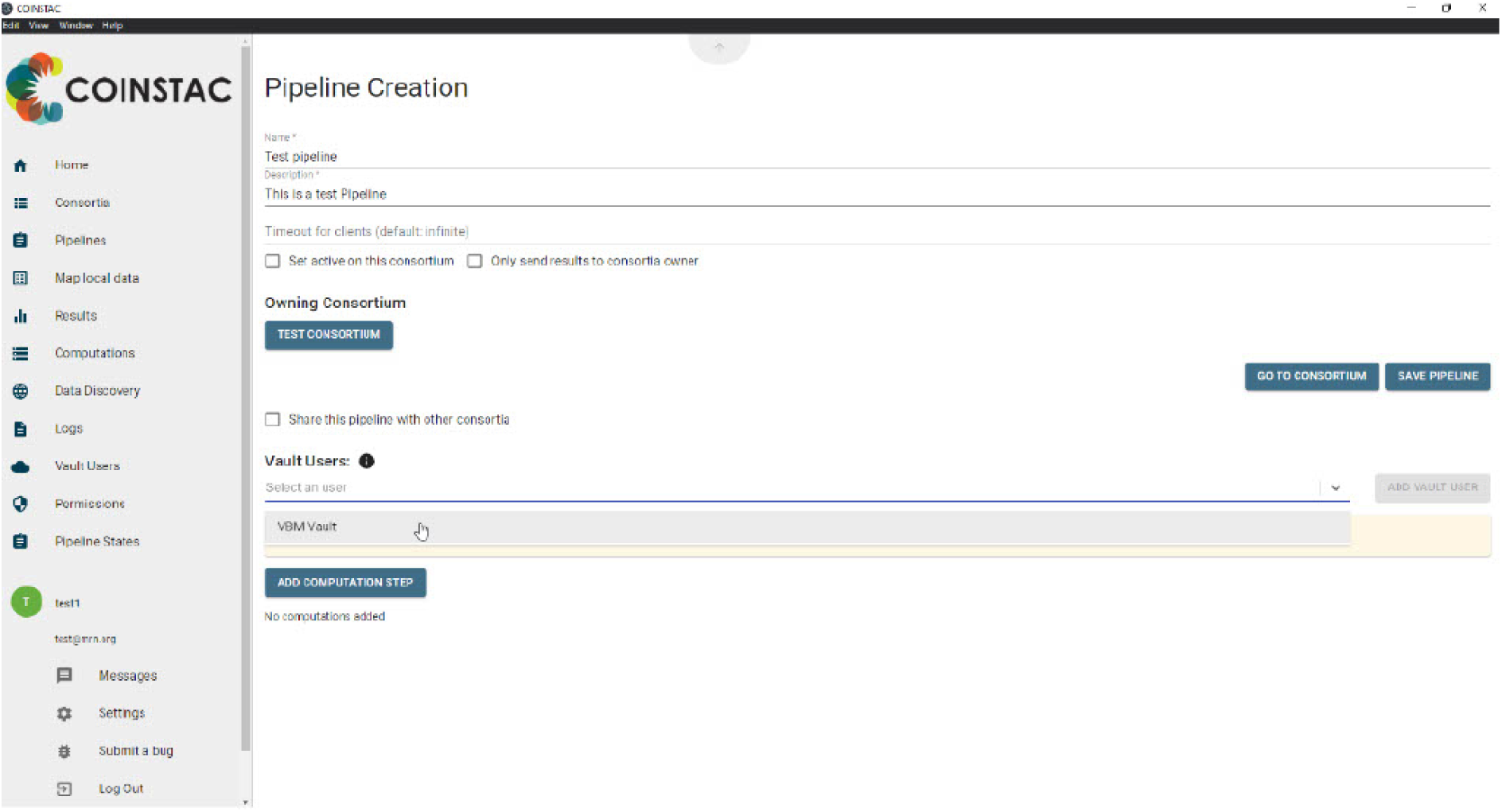
Adding Vault data to an analysis pipeline

#### 2.2.3 Hosting Vaults

Making datasets available for federated analysis through COINSTAC is simple using Vaults. Vaults can be hosted in a variety of compute environments such as: on personal machines, on-premises servers, on a cluster of compute nodes, or in a virtual cloud. Both publicly available datasets and private datasets can be made available to the COINSTAC platform via Vaults. COINSTAC consortia can include any combination of diverse types of data: public and private datasets, data hosted on local machines, TReNDS-hosted Vaults, and third-party Vaults connected to COINSTAC as shown in Fig. 2.

**Figure 2.**
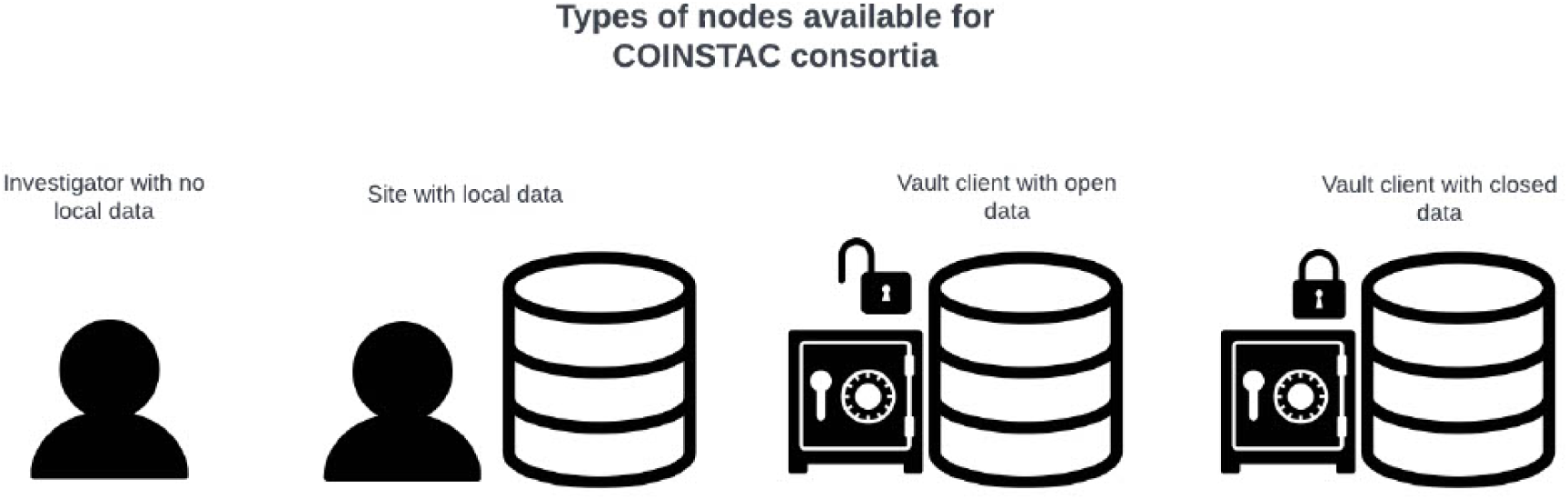
Different types of participants interacting with COINSTAC

In addition to TReNDS-hosted vaults, data owners are able to host their own (public or private) data as Vaults (Fig. 3) by using the coinstac-vault-client software package at https://www.npmjs.com/ `package/coinstac-vault-client`.

**Figure 3.**
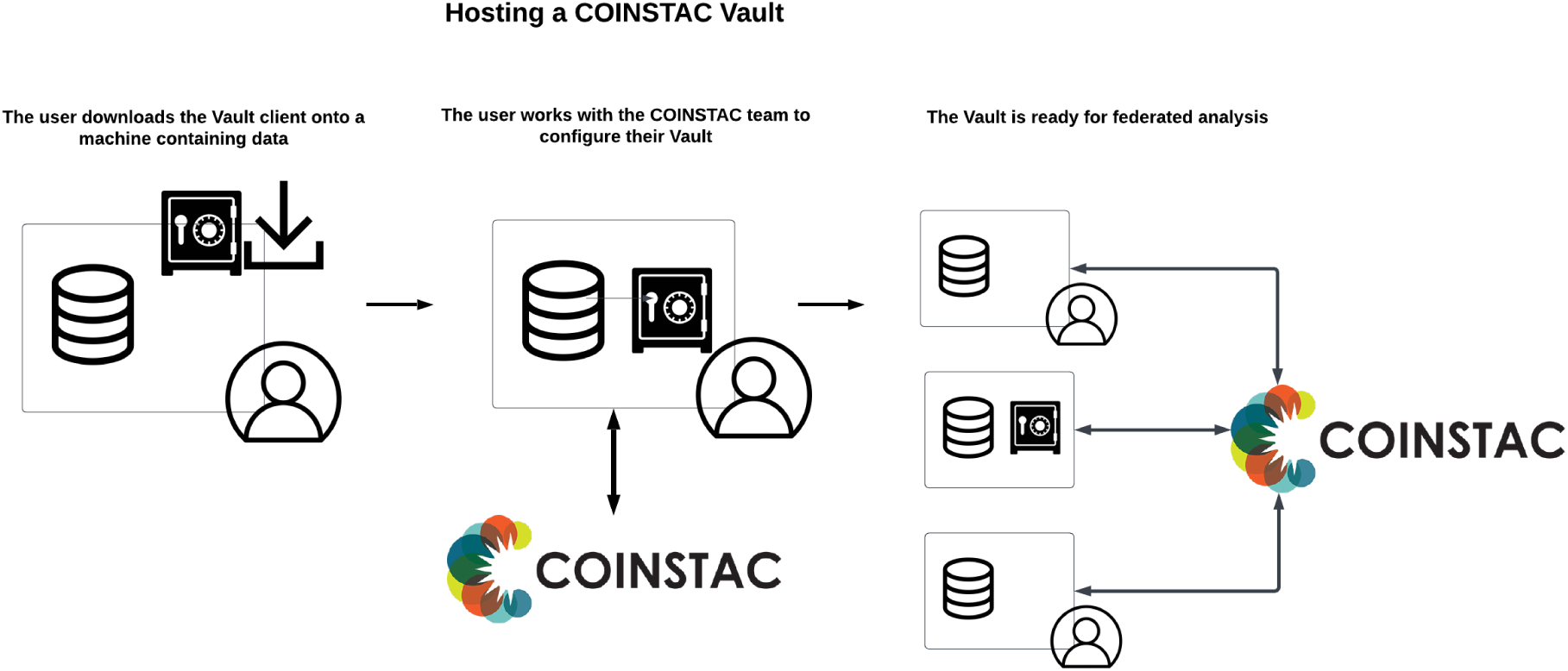
Process of creating a Vault in COINSTAC

The process for hosting a dataset in a Vault is described below:

- The user installs the Vault client on the host machine.
- The user configures the Vault client by specifying the directory of the dataset.
- The user makes a request to the COINSTAC team to have the Vault be added to the COINSTAC ecosystem.
- The COINSTAC team will give the user API keys to be added to the user’s local Vault configuration.

After this process, the Vault becomes available for use in the COINSTAC system. Consortium owners can select to include the Vault in their consortium and perform federated analysis using Vault data. Whether the data was downloaded from a public repository or collected privately, the process is the same for both types of data since the source data stays on the user’s local machine.

#### 2.2.4 Vault architecture overview

The Vault client software package is built upon the same core code as the COINSTAC desktop application to manage containers and execute computation pipelines. However, it omits the user interface (UI) component and includes additional code that enables the client to be persistently online and available. The desktop application has been modified to allow consortium owners to add Vault clients to their consortium via the GUI.

The Vault client is a NodeJS server running on the local machine, responsible for maintaining a persistent connection with the COINSTAC system using the coinstac-vault-client package. The server communicates with the COINSTAC central server using websockets and HTTP protocols. It manages the life-cycle of containers (Docker, Singularity) through the coinstac-container-manager package, which is responsible for isolating and executing the computations within the federated analyses. The Vault client also utilizes other core COINSTAC libraries such as coinstac-client-core, coinstac-client-server, coinstac-pipeline, and coinstac-common, all of which are npm packages, to ensure seamless integration with the COINSTAC ecosystem. An overview is shown in figure 4.

**Figure 4.**
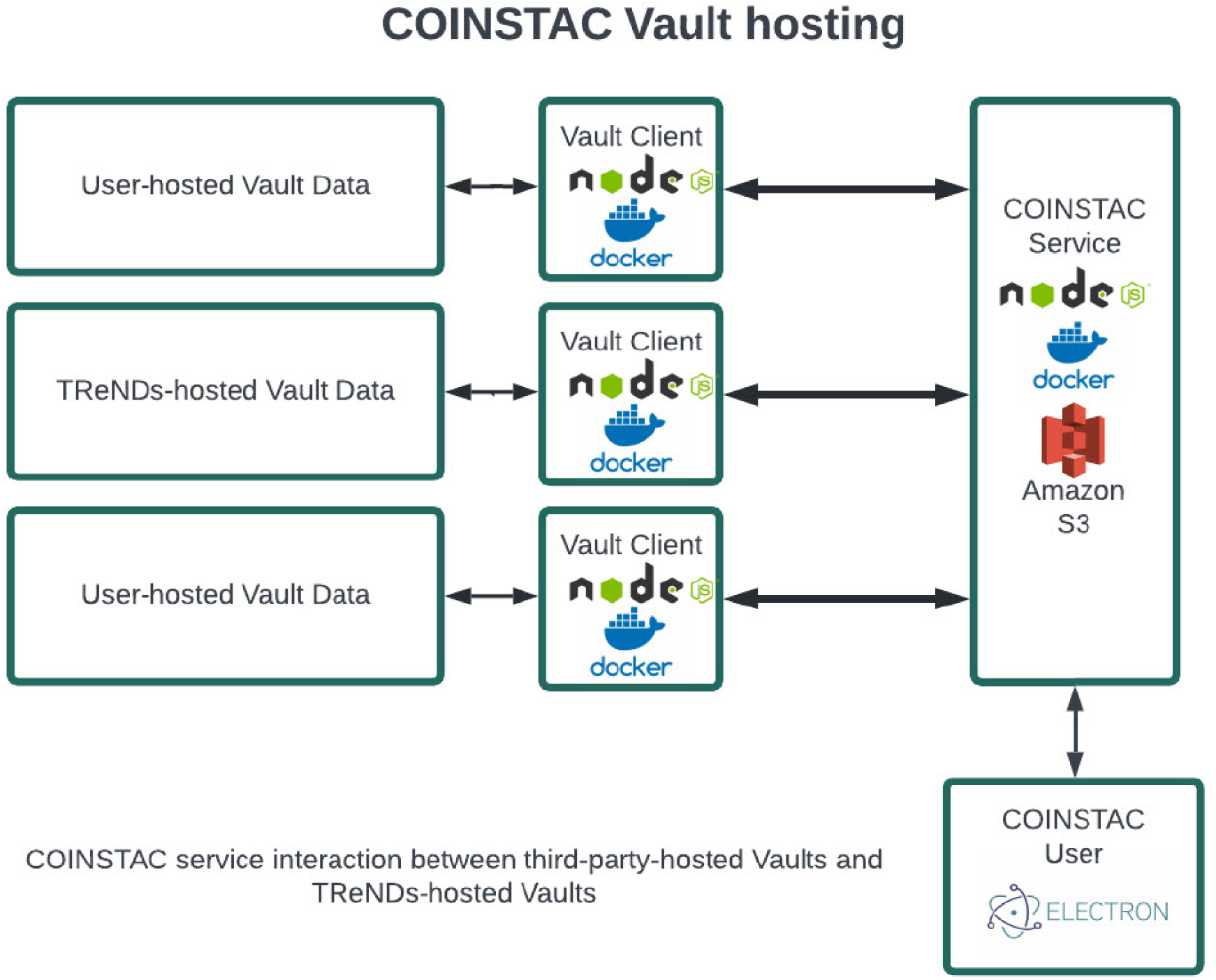
Architecture of Vaults in COINSTAC

Message passing, which is an integral part of federated analyses, is handled by the Vault client using MQTT (MQ Telemetry Transport) and HTTP protocols. MQTT is a lightweight messaging protocol optimized for high-latency or unreliable networks.

For pipeline runs in consortia that only use Vaults, the result data is uploaded to a secure Amazon S3 bucket, which can then be downloaded by consortium members using the desktop application. This ensures that the results are securely stored and easily accessible by authorized users.

In summary, the Vault architecture in COINSTAC improves the overall efficiency and user experience of performing federated analyses. By maintaining a persistent connection, the Vault client ensures that datasets are readily available for analysis without the need for manual intervention by data owners. Additionally, the integration of the Vault client within the COINSTAC ecosystem allows for seamless interaction between the desktop application and the Vaults, making it simple for consortium owners to include Vault data in their federated analyses.

#### 2.2.5 Vault use-cases

In this section, we present various use-cases that highlight the benefits and versatility of Vaults in COINSTAC.

##### 2.2.5.1 Curated Vaults

TReNDS actively curates and hosts public datasets, making them readily available for the COINSTAC community through the creation of Vaults. These curated Vaults ensure that the public datasets are vetted, of high quality, and easily accessible. Users can contribute to this initiative by hosting Vaults for other public datasets, further expanding the range of data resources available within COINSTAC.

##### 2.2.5.2 User with local data

A researcher with a local dataset can benefit from Vaults by accessing and combining data from additional datasets containing relevant variables. This process is particularly useful when the researcher’s local data is insufficient for conducting a robust analysis. By collaborating with other COINSTAC consortium members and incorporating data from Vaults, the researcher can increase the sample size and statistical power of their study without requiring a Data Use Agreement (DUA) or other additional steps.

##### 2.2.5.3 User with no local data

For investigators who do not have their own data but want to analyze existing datasets, Vaults provide a valuable solution. The investigator can create a consortium, add selected Vaults using the COINSTAC UI, and initiate the analysis. This approach enables the investigator to obtain meaningful insights from existing datasets without needing to coordinate with the Vault data owners.

##### 2.2.5.4 User with limited storage/computing resources

Vaults are also advantageous for researchers with limited storage or computing resources. For example, a researcher with a low-powered laptop and minimal storage capacity can still analyze large datasets by creating a consortium and running an analysis using only Vault clients. The data processing occurs on the respective Vault servers, and the results are sent back to the investigator, eliminating the need for high-capacity local hardware.

By addressing these diverse use-cases, COINSTAC Vaults offer a flexible and efficient solution for researchers to access, collaborate, and analyze datasets in a federated environment.

## 3 RESULTS

In this section, we provide details of several different types of Vaults hosted on COINSTAC and also perform some analysis using these Vaults showing that adding of Vault data increases sample size and statistical power of results.

### 3.1 TReNDS VBM COBRE

The TReNDS VBM COBRE Vault contains structural MRI images from 152 participants, approximately half healthy volunteers and half individuals diagnosed with schizophrenia, collected as part of the Mind Research Network COBRE study Aine et al. (2017). The Vault includes gray matter MRI images that have been run through a VBM preprocessing pipeline in the SPM toolbox. In addition, we have demographic information, symptom severity scales, and cognitive measures to select from when building a desired model. Figure 5 shows the beta images from running VBM regression on all the voxels from normalized smoothed gray matter images from the TReNDS COBRE Vault. Age, sex, and diagnosis information were used as covariates in the regression model. Results show decreases in brain volume with age, reduced volume in visual areas and along the gray/white boundary in females, and reduced volume in insular-temporal and medial frontal regions in schizophrenia patients, consistent with previous results.

**Figure 5.**
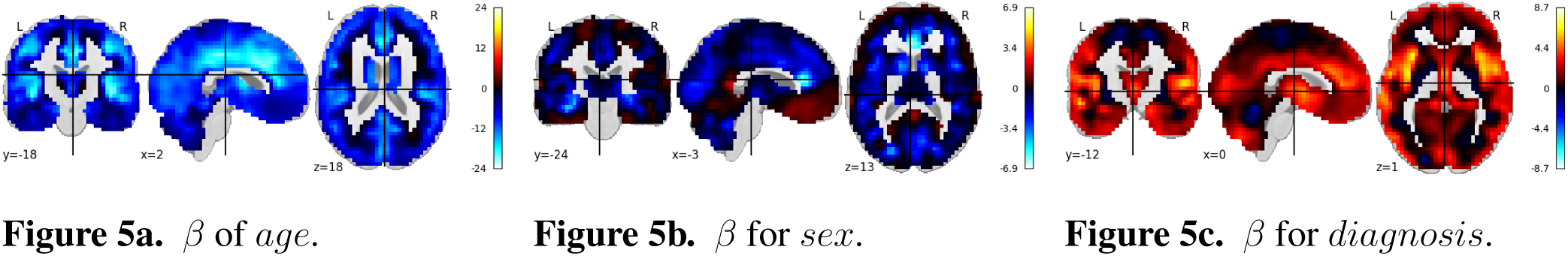
Rendered images show voxel-wise *β* values corresponding to the *age*, *sex* and *diagnosis* covariates using COBRE VBM data Vault in COINSTAC. For age, negative values show that the gray matter volume decreases with age. For sex, positive values indicate male’s gray matter volume is greater than female’s gray matter volume and vice versa. For Diagnosis, positive values indicate control’s gray matter volume is greater than patient’s gray matter volume and vice versa.

The following section describes this use-case with 55 participant’s structural MRI scans collected under MCIC project Gollub (2013). The results from running regression on the normalized smoothed gray matter images from this project are shown in figure 6.

**Figure 6.**
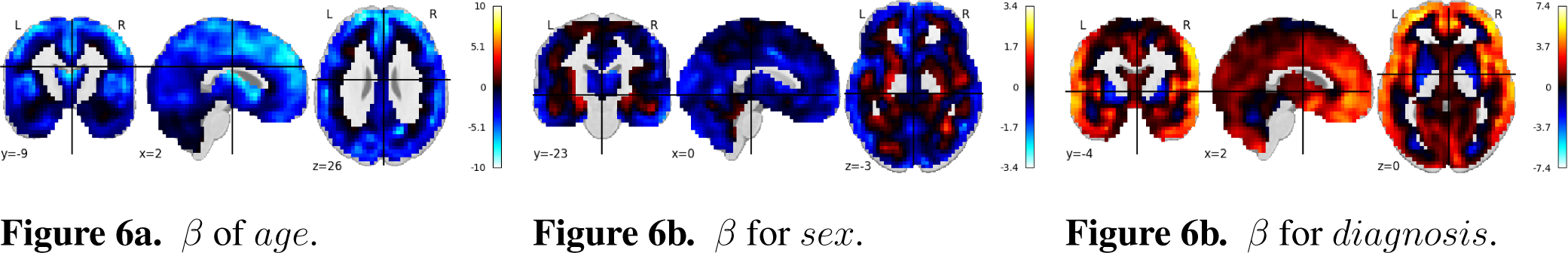
Rendered images show voxel-wise *β* values corresponding to the *age*, *sex*, and *diagnosis* covariates using MCIC *s*MRI data in COINSTAC. For age, negative values show that the gray matter volume decreases with age. For sex, positive values indicate male’s gray matter volume is greater than female’s gray matter volume and vice versa. For Diagnosis, positive values indicate control’s gray matter volume is greater than patient’s gray matter volume and vice versa.

Using the MCIC dataset, we similarly see widespread reduction in brain volume for age, visual and gray/white boundary reductions in volume in females, and insular-temporal and medial frontal (as well as more wide spread) reductions in schizophrenia patients.

The TReNDS VBM COBRE Vault was combined with the MCIC dataset, allowing for an increased sample size, in the same regression analysis to examine diagnostic effects while accounting for age and sex. The combined dataset was largely consistent with the individual site analysis, with the exception of the male/female effect which shows a more complex pattern of increases and decreases, though still largely conforming to reductions in white/gray matter boundary and primary visual area volumes. Gupta et al. (2015). Results of this study are shown in figure 7.

**Figure 7.**
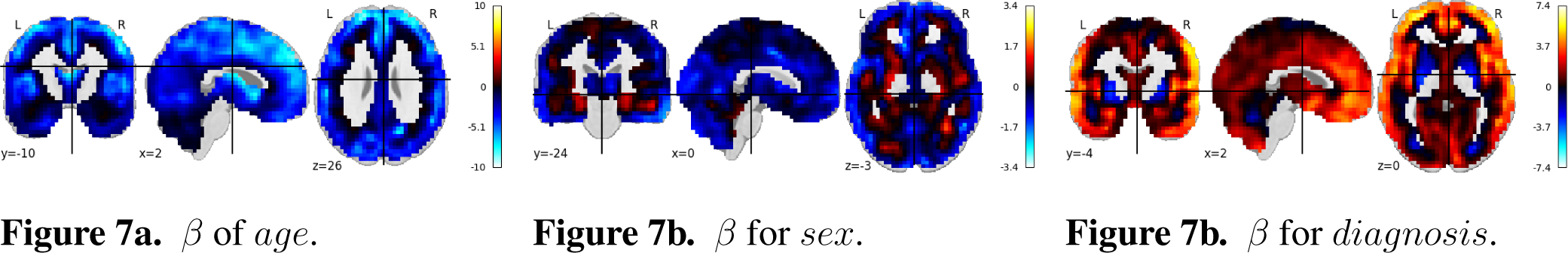
MCIC+COBRE Vault: Rendered images show voxel-wise *β* values corresponding to the *age*, *sex*, and *diagnosis* covariates using MCIC *s*MRI data along with the data in the COBRE Vault in COINSTAC. For age, negative values show that the gray matter volume decreases with age. For sex, positive values indicate male’s gray matter volume is greater than female’s gray matter volume and vice versa. For Diagnosis, positive values indicate control’s gray matter volume is greater than patient’s gray matter volume and vice versa.

### 3.2 TReNDS FreeSurfer COBRE

This Vault contains data from 152 subjects, approximately half controls and half individuals with chronic schizophrenia, collected as part of the Mind Research Network COBRE study ^7^. The Vault includes cortical and sub-cortical volumetric and surface-based measurements from two FreeSurfer atlases, Desikan-Killiany and Destrieux. In addition, we have a total of 11 variables across demographic, cognitive and substance use to select from when building a desired model.

We ran Ridge regression on the above Vault data on Freesurfer volumetric and surface based measurements on about 500 regions of interest. We noticed the following differences between controls and patients.

Controls have higher values in temporal lobe, as shown in the thickness measurements of tables 1 2 3 4 5

**Table 1.** Global freesurfer stats for *lh S temporal inf thickness*

| Global stats – lh_S_temporal_inf_thickness | $\beta_0$ (const) | $\beta_1$ (age) | $\beta_2$ (sex) | $\beta_3$ (isControl.True) |
| --- | --- | --- | --- | --- |
| Coefficient | 2.5552 | -0.0052 | 0.0323 | 0.1071 |
| t Stat | 44.8963 | -5.014 | 1.0642 | 4.1085 |
| P-value | 0 | 0 | 0.289 | 1.00E-04 |
| R Squared | 0.237444911 |  |  |  |
| Degrees of Freedom | 145 |  |  |  |

**Table 2.** Global freesurfer stats for *rh S oc − temp lat thickness*

| Global Stats – rh_S_oc-temp_lat_thickness | $\beta_0$ (const) | $\beta_1$ (age) | $\beta_2$ (sex) | $\beta_3$ (isControl.True) |
| --- | --- | --- | --- | --- |
| Coefficient | 2.5331 | -0.0036 | 0.0127 | 0.1158 |
| t Stat | 38.8572 | -3.0471 | 0.3663 | 3.878 |
| P-value | 0 | 0.0027 | 0.7147 | 2.00E-04 |
| R Squared | 0.149884354 |  |  |  |
| Degrees of Freedom | 145 |  |  |  |

**Table 3.** Global freesurfer stats for *lh middletemporal thickness*

|  |  |  |  |  |
| --- | --- | --- | --- | --- |
| Global Stats – lh_middletemporal_thickness | $\beta 0$ (const) | $\beta 1$ (age) | $\beta 2$ (sex) | $\beta 3$ (isControl_True) |
| Coefficient | 3.0038 | -0.0057 | -0.0134 | 0.0829 |
| t Stat | 60.2257 | -6.3161 | -0.5056 | 3.63 |
| P-value | 0 | 0 | 0.6139 | 4.00E-04 |
| R Squared | 0.275552216 |  |  |  |
| Degrees of Freedom | 145 |  |  |  |

**Table 4.** Global freesurfer stats for *lh superiortemporal thickness*

|  |  |  |  |  |
| --- | --- | --- | --- | --- |
| Global Stats – lh_superiortemporal_thickness | $\beta 0$ (const) | $\beta 1$ (age) | $\beta 2$ (sex) | $\beta 3$ (isControl_True) |
| Coefficient | 2.9341 | -0.0067 | 0.0169 | 0.0682 |
| t Stat | 52.568 | -6.6199 | 0.5679 | 2.6659 |
| P-value | 0 | 0 | 0.571 | 0.0085 |
| R Squared | 0.266849477 |  |  |  |
| Degrees of Freedom | 145 |  |  |  |

**Table 5.**
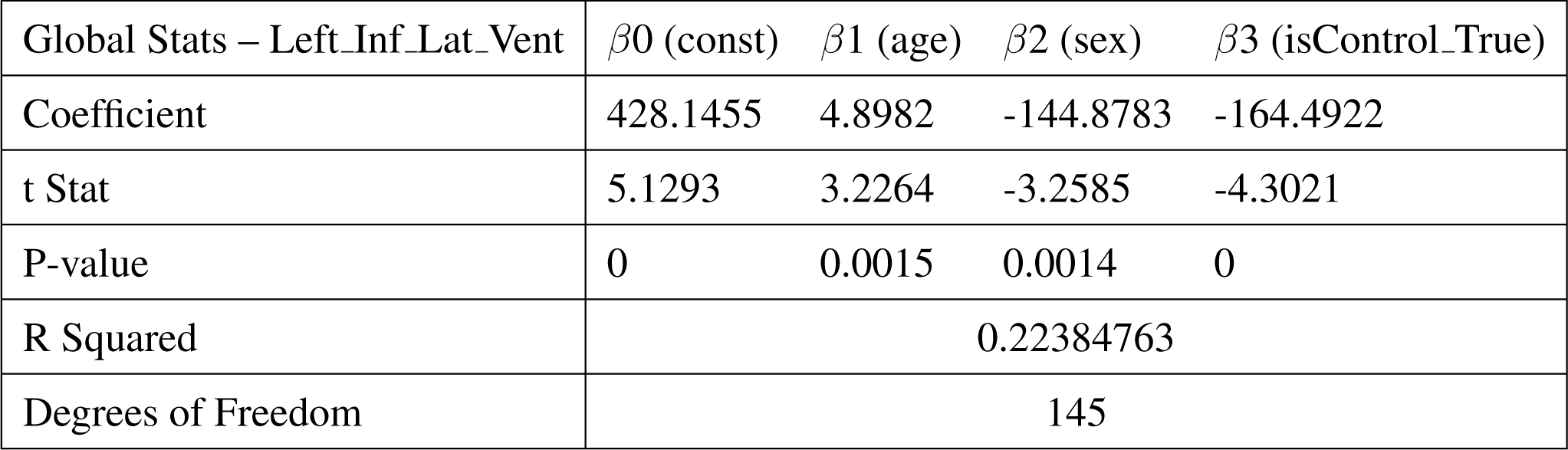
Global freesurfer stats for *Left Inf Lat V ent*

### 3.3 Child Mind Institute (CMI) VBM VAULT

This Vault contains data from 922 children and adolescents (ages 6 – 22, 603 Male and 319 female), collected as part of the Healthy Brain Network study Alexander et al. (2017). The Vault includes gray matter segmentation data from an SPM VBM preprocessing pipeline. In addition, we have a total of 11 variables across various demographic, cognitive and substance use domains to select from when building a desired model.

Figure 8 shows the beta images from running regression on all the voxels from normalized smoothed gray matter images from the CMI VBM VAULT. Age and sex were used as covariates in the regression model. Results were largely consistent with those from the MCIC and COBRE analyses, showing widespread volume reductions with age, and reductions along the gray/white matter boundary in females.

**Figure 8.**
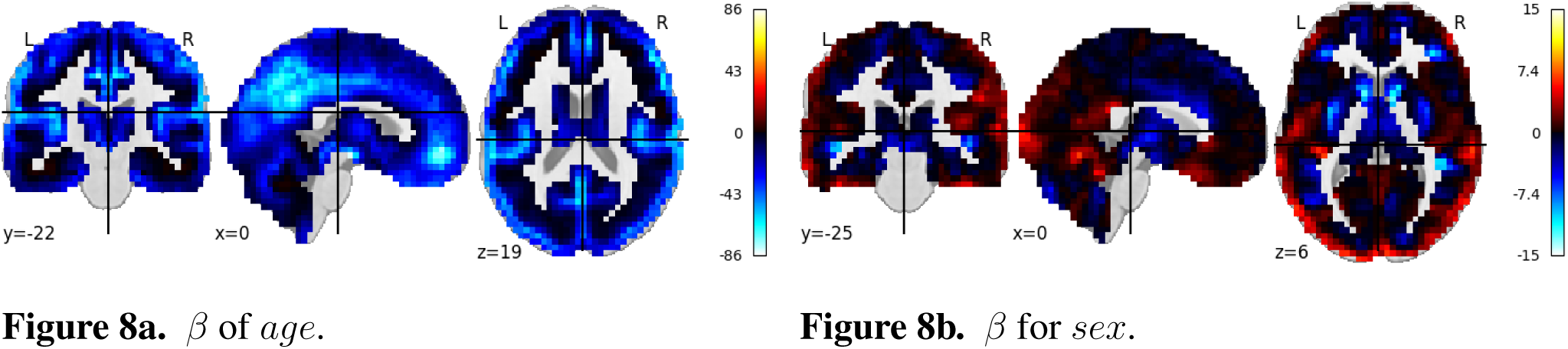
Rendered images show voxel-wise *β* values corresponding to the *age* and *sex* covariates using CMI VBM Vault data in COINSTAC. For age, negative values show that the gray matter volume decreases with age. For sex, positive values indicate male’s gray matter volume is greater than female’s gray matter volume and vice versa.

### 3.4 TReNDS NeuroMark Group-ICA COBRE VAULT

Group ICA Calhoun et al. (2001) is one of the frequently used preprocessing computations for neuroimaging data. Data preprocessed with group ICA can be used to perform different types of analyses. This GICA Vault comprises data from 189 subjects from the COBRE project analyzed with Neuromark template which uses 66 predefined ROIs. This Vault data includes independent component analysis (ICA) maps, Functional network connectivity maps (FNC) data etc. that have been generated using spatially constrained ICA with the `Neuromark fMRI 1.0 template`(available in the GIFT software^8^,^9^) including 53 intrinsic networks (components). This Vault data can be readily used for secondary analysis like mancova. In this case, we use the GICA pre-processed data from the Vault to perform univariate regression analysis, the results of which are shown in figure 9.

**Figure 9.**
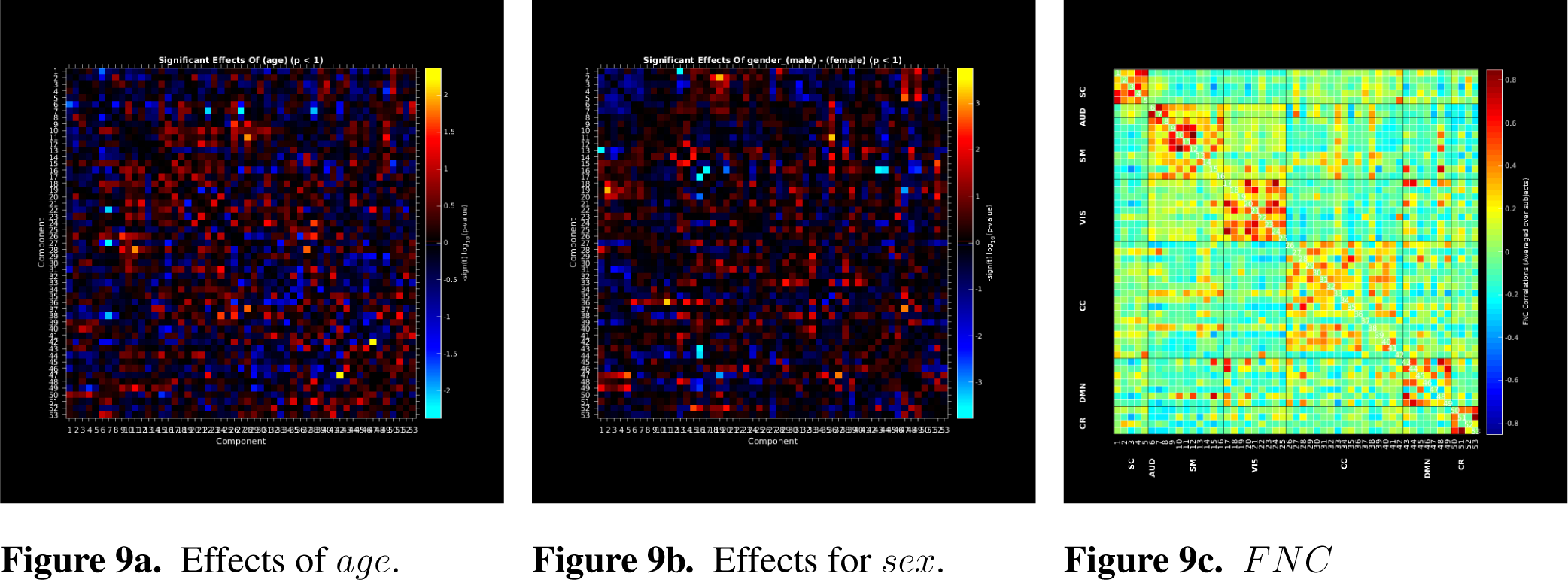
Rendered images show univariate regression results demonstrating the effects of age and sex on correlation between the 53 independent components and FNC correlation map using Vault data in COINSTAC.

**Figure 10.**
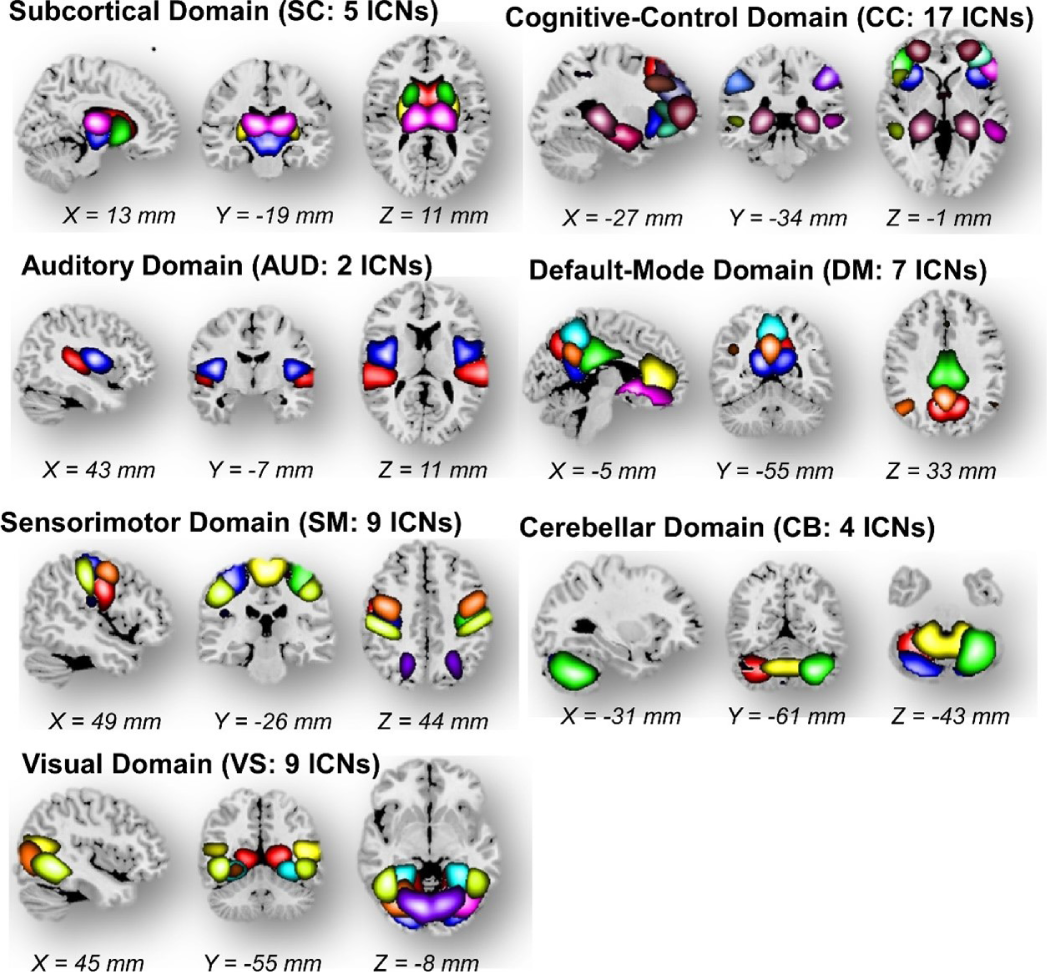
The Neuromark fMRI 1.0 template with 53 intrinsic networks (components) from 7 major networks.

The Neuromark fMRI domains identified in Du et al. Briefly, these seven identified network templates were divided based on anatomical and functional propertiesDu et al. (2020). In each subfigures, one color in the composite maps corresponds to an intrinsic connectivity network (ICN). The Neuromark fMRI 1.0 template is available in the GIFT software10.

## 4 DISCUSSION

In recent decades, data sharing has driven substantial advancements in the field of neuroimaging and expanded opportunities for open science collaboration. Although data sharing has undeniable merits, it also faces inherent limitations, including technological, policy, administrative, and methodological barriers that can hinder progress. COINSTAC Vaults and the federated computing framework within COINSTAC uniquely address these challenges by enabling data analysis while maintaining privacy protection, specifically in the context of neuroimaging research. The ‘always-on’ status of Vaults streamlines collaboration between institutions by eliminating the need for synchronized efforts across users. The accessibility and user-friendly interface of COINSTAC Vaults serve as powerful tools for reproducible research, an area that has faced significant criticism in recent years. By bolstering the collaborative capabilities of federated learning and addressing the limitations of traditional data sharing, COINSTAC Vaults provide a cutting-edge solution for the neuroimaging community, pushing the boundaries of data analysis and open science.

COINSTAC offers a user-friendly GUI for the neuroimaging field, enabling federated learning on neuroimaging data with ease. Its extensive library includes numerous algorithms and pipelines, facilitating efficient processing of large datasets. Currently, over twenty computations are available in open-source repositories, allowing users to create versatile analytic pipelines. The integration of Vaults further enhances the user experience by providing access to diverse datasets, enabling efficient analysis with robust data, and fostering collaboration across institutions asynchronously.

Compared to OpenNeuro ^10^, and OpenfMRI A.Poldrack and Gorgolewski (2017) like projects, where users can access data, download them and perform analysis on their own, Vaults allow users to perform neuroimaging analysis in federated learning platform immediately, without the need to download data and toolboxes onto a centralized computing environment. Vaults can help researchers to run an initial test on a data or their algorithm quickly to help setup their hypotheses or validate it to save time before they commit to a big project.

In addition to being faster to execute by being immediately available with no downloading or manual coordination, curated Vaults that follow documented standards make studies easier to design, execute, and reproduce. For example: Neuroimaging datasets can contain a large number of variables that apply to each subject: demographic information, cognitive measures, etc. The number of these variables can range from tens to hundreds. Using standard naming conventions makes it easier for researchers to understand what each variable tracks so that they can select the relevant variables for their study. Standard and predictable ways for handling missing data in Vaults makes it easier for researchers to design their analyses.

COINSTAC is unique in its commitment to open science, with its open-source platform promoting seamless integration of modular computations and streamlining federated analyses. The addition of COINSTAC Vaults reinforces this commitment by simplifying dataset inclusion in federated analyses, encouraging community contributions, and preserving privacy for private datasets. By offering easy access to public datasets and enabling secure contributions from private dataset owners, COINSTAC Vaults foster collaboration and dedication to open science.

### 4.1 Limitations and Challenges

COINSTAC Vaults offer numerous benefits, but there are also limitations and challenges to consider. One concern is that allowing arbitrary summary queries on a dataset might enable an attacker to reconstruct the data. To mitigate such risks, the system must be privacy-preserving from “end-to-end,” incorporating techniques like secure multiparty computation or differential privacy. Implementing these methods can be difficult due to floating point implementation issues Ilvento (2020a,b); Mironov (2012) and the introduction of noise, which may increase error or variance in the analysis results.

While differentially private algorithms can provide stronger privacy guarantees, sharing data derivatives without differential privacy might be adequate in some situations, depending on the trust model and privacy concerns of data holders. These issues should be addressed on a case-by-case basis.

Additional challenges include data distribution, network bandwidth, and communication speed. Federated learning and open-source solutions can help address some of these problems, but further research and development are needed to optimize COINSTAC Vaults’ performance in various research settings. Our “Decentralized Sparse Deep Artificial Neural Networks in COINSTAC (CPU and GPU enabled)” algorithm allows users to save network bandwidth when transferring thousands of derived data/machine learning parameters across nodes.

In summary, COINSTAC Vaults mark a significant advancement in federated neuroimaging research, data privacy preservation, and open science promotion. By tackling the existing limitations and challenges, COINSTAC Vaults can further improve collaboration and innovation within the field.

## 5 CONCLUSION

The neuroimaging field is experiencing rapid growth, generating substantial data volumes. However, access to this data is challenged by technological, privacy, administrative, and methodological constraints. In this study, we present COINSTAC Vaults as a solution that streamlines data access and analysis, specifically in the context of neuroimaging research. COINSTAC Vaults ensure continuous availability of high-quality data, promoting the advancement of open science and fostering efficient collaboration between researchers.

We invite researchers to use COINSTAC Vaults in their studies and to host their own datasets using COINSTAC Vaults. By adopting COINSTAC Vaults, the neuroimaging community can overcome the barriers associated with traditional data sharing and analysis methods, paving the way for groundbreaking discoveries.

### 5.1 Future Work

The long-term vision for COINSTAC and COINSTAC Vaults includes:

- Introducing new user interface features, such as the ability to search Vaults and filter by covariates, to improve user experience and efficiency.
- Making new datasets available as Vaults, including those from OpenNeuro, the Autism Brain Imaging Data Exchange (ABIDE), the National Institute of Mental Health Data Archive (NDA), the Open Access Series of Imaging Studies (OASIS), and the Image and Data Archive (IDA), to enhance the diversity of Vaults.
- Increase BIDS (Brain Imaging Data Structure) support to all major neuroimaging modalities and Vault datasets, to ensure interoperability and ease of use.
- Increase compliance to programs such as the FAIR (Findability, Accessibility, Interoperability, and Reuse) Guiding Principles for scientific data management and stewardship, to enhance the overall data sharing ecosystem.
- Exploring the integration of differential privacy techniques to further safeguard data privacy, while preserving the utility of data analysis.

## Data availability statement

Generated Statement: Publicly available datasets were analyzed in this study. This data can be found here: http://fcon_1000.projects.nitrc.org/indi/retro/cobre.html.

## AUTHOR CONTRIBUTIONS

DM, RK, VDC: conceptualization. DM, SB, SaPa, and PP: methodology. DM, SB, SaPa, KRM, PP, BB, JR: writing—original draft preparation. SB, SaPa: data analysis. SP and VDC: supervision. All authors writing—review and editing, read, and agreed to the published version of the manuscript.

## Contribution to the field

Neuroimaging research faces numerous challenges, including technological, policy, administrative, and methodological barriers that hinder collaboration and data sharing. COINSTAC is an already established open-source platform that advances open science by streamlining federated analyses and facilitating seamless integration of modular computations. This study presents the new COINSTAC Vaults (CVs), an enhancement to the COINSTAC platform, which addresses the current challenges of the field by offering standardized, persistent, and highly-available datasets, integrating seamlessly with federated analysis capabilities. In this paper, we demonstrate the impact of CVs through several studies utilizing federated analysis, showcasing their potential to improve the reproducibility of research and increase sample sizes in neuroimaging studies. COINSTAC Vaults improve the reproducibility of research and enable larger sample sizes in neuroimaging studies. Unlike traditional data sharing platforms, CVs facilitate federated learning, ensuring privacy protection and fostering asynchronous collaboration between institutions. By offering easy access to public datasets and secure contributions from private dataset owners, CVs promote open science and data-driven discovery.

## Ethics statements

### Studies involving animal subjects

Generated Statement: No animal studies are presented in this manuscript.

### Studies involving human subjects

Generated Statement: No human studies are presented in this manuscript.

### Inclusion of identifiable human data

Generated Statement: No potentially identifiable human images or data is presented in this study.

## Footnotes

1 https://nda.nih.gov/

2 https://www.speedtest.net/global-index

3 https://www.apple.com/macbook-pro-14-and-16/specs/

4 http://coinstac.trendscenter.org

5 https://coinstac.org/

6 https://github.com/trendscenter/coinstac

7 http://fcon_1000.projects.nitrc.org/indi/retro/cobre.html

8 http://trendscenter.org/software/gift

9 http://trendscenter.org/data

10 https://openneuro.org/

## REFERENCES

Aine, C. J., Bockholt, H. J., Bustillo, J. R., Cañive, J. M., Caprihan, A., Gasparovic, C., et al. (2017). Multimodal neuroimaging in schizophrenia: Description and dissemination. Neuroinformatics 15, 343–364. doi:10.1007/s12021-017-9338-9

Alexander, L. M., Escalera, J., Ai, L., Andreotti, C., Febre, K., Mangone, A., et al. (2017). An open resource for transdiagnostic research in pediatric mental health and learning disorders. Scientific Data 4, 1–26. doi:10.1038/sdata.2017.181

Andrade, C. (2020). Sample size and its importance in research. Indian Journal of Psychological Medicine 42, 102–103. doi:10.4103/IJPSYM.IJPSYM\_504\_19

A. Poldrack, R. and Gorgolewski, K. J. (2017). OpenfMRI: Open sharing of task fMRI data. NeuroImage 144 part B. doi:10.1016/j.neuroimage.2015.05.073

[Dataset] Babayan, A., Baczkowski, B., Cozatland, R., Dreyer, M., Engen, H., Erbey, M., et al. (2020-07-22). MPI-Leipzig Mind-Brain-Body Dataset

Biswal, B. B., Mennes, M., Zuo, X.-N., and Milham, M. P. (2010). Toward discovery science of human brain function. Proceedings of the National Academy of Sciences 107, 4734–4739

Bonawitz, K., Ivanov, V., Kreuter, B., Marcedone, A., McMahan, H. B., Patel, S., et al. (2016). Practical Secure Aggregation for Federated Learning on User-Held Data. Tech. Rep. arXiv:1611.04482 [cs.CR], ArXiV

Bonawitz, K., Ivanov, V., Kreuter, B., Marcedone, A., McMahan, H. B., Patel, S., et al. (2017). Practical Secure Aggregation for Privacy-Preserving Machine Learning. In Proceedings of the 2017 ACM SIGSAC Conference on Computer and Communications Security (New York, NY, USA: ACM), CCS ’17, 1175–1191. doi:10.1145/3133956.3133982

Calhoun, V. D., Adali, T., Pearlson, G., and Pekar, J. (2001). Group ICA of functional MRI data: separability, stationarity, and inference. In Proceeedings of the International Conference on ICA and BSS. 155

Du, Y., Fu, Z., Sui, J., Gao, S., Xing, Y., Lin, D., et al. (2020). NeuroMark: An automated and adaptive ICA based pipeline to identify reproducible fMRI markers of brain disorders. NeuroImage: Clinical 28, 102375. doi:10.1016/j.nicl.2020.102375. Epub 2020 Aug 11

Dwork, C. and Roth, A. (2013). The algorithmic foundations of differential privacy. Foundations and Trends in Theoretical Computer Science 9, 211–407. doi:10.1561/0400000042

Esteban, O., Markiewicz, C. J., Blair, R. W., Moodie, C. A., Isik, A. I., Erramuzpe, A., et al. (2019). fMRIPrep: a robust preprocessing pipeline for functional MRI. Nature Methods 16, 111–116. doi:10.1038/s41592-018-0235-4

Gazula, H., Kelly, R., Romero, J., Verner, E., Baker, B. T., Silva, R. F., et al. (2020). COINSTAC: Collaborative informatics and neuroimaging suite toolkit for anonymous computation. Journal of Open Source Software 5, 2166. doi:10.21105/joss.02166

Gazula, H., Rootes-Murdy, K., Holla, B., Basodi, S., Zhang, Z., Verner, E., et al. (2023). Federated analysis in COINSTAC reveals functional network connectivity and spectral links to smoking and alcohol consumption in nearly 2,000 adolescent brains. Neuroinformatics 21, 287–301. doi:10.1007/s12021-022-09604-4

Gollub, R. L. (2013). The MCIC collection: a shared repository of multi-modal, multi-site brain image data from a clinical investigation of schizophrenia. Neuroinformatics 11, 367–388. doi:10.1007/s12021-013-9184-3

Gorgolewski, K. J., Auer, T., Calhoun, V. D., Craddock, R. C., Das, S., Duff, E. P., et al. (2016). The brain imaging data structure, a format for organizing and describing outputs of neuroimaging experiments. Scientific Data 3, 1–9. doi:10.1038/sdata.2016.44

Gupta, C. N., Calhoun, V. D., Rachakonda, S., Chen, J., Patel, V., Liu, J., et al. (2015). Patterns of gray matter abnormalities in schizophrenia based on an international mega-analysis. Schizophrenia Bulletin 41, 1133–1142. doi:10.1093/schbul/sbu177

Heikkilä, M. A., Koskela, A., Shimizu, K., Kaski, S., and Honkela, A. (2020). Differentially private cross-silo federated learning. Tech. Rep. arXiv:2007.05553 [cs.CR], ArXiV

Homer, N., Szelinger, S., Redman, M., Duggan, D., Tembe, W., Muehling, J., et al. (2008). Resolving individuals contributing trace amounts of dna to highly complex mixtures using high-density snp genotyping microarrays. PLoS genetics 4, e1000167

Ilvento, C. (2020a). Implementing differentially private integer partitions. Presented at the 2020 Workshop on the Theory and Practice of Differential Privacy

Ilvento, C. (2020b). Implementing sparse vector. Presented at the 2020 Workshop on the Theory and Practice of Differential Privacy

Imtiaz, H., Mohammadi, J., Silva, R., Baker, B., Plis, S. M., Sarwate, A. D., et al. (2021). A correlated noise-assisted decentralized differentially private estimation protocol, and its application to fMRI source separation. IEEE Transactions on Signal Processing 69, 6355–6370. doi:10.1109/TSP.2021.3126546

Jwa, A. S. and Poldrack, R. A. (2022). The spectrum of data sharing policies in neuroimaging data repositories. Human Brain Mapping 43. doi:10.1002/hbm.25803

Kairouz, P., McMahan, H. B., Avent, B., Bellet, A., Bennis, M., Bhagoji, A. N., et al. (2021). Advances and open problems in federated learning. Foundations and Trends® in Machine Learning 14, 1–210. doi:10.1561/2200000083

Markiewicz, C. J., Gorgolewski, K. J., Feingold, F., Blair, R., Halchenko, Y. O., Miller, E., et al. (2021). The OpenNeuro resource for sharing of neuroscience data. eLife 10, e71774. doi:10.7554/eLife.71774

McGuire, A. L., Basford, M., Dressler, L. G., Fullerton, S. M., Koenig, B. A., Li, R., et al. (2011). Ethical and practical challenges of sharing data from genome-wide association studies: the emerge consortium experience. Genome research 21, 1001–1007

Ming, J., Verner, E., Sarwate, A., Kelly, R., Reed, C., Kahleck, T., et al. (2017). COINSTAC: Decentralizing the future of brain imaging analysis. F1000Research 6, 1512. doi:10.12688/f1000research.12353.1

Mironov, I. (2012). On significance of the least significant bits for differential privacy. In Proceedings of the 2012 ACM Conference on Computer and Communications Security (CCS). 650–661. doi:10.1145/2382196.2382264

Plis, S. M., Sarwate, A. D., Wood, D., Dieringer, C., Landis, D., Reed, C., et al. (2016). COINSTAC: A privacy enabled model and prototype for leveraging and processing decentralized brain imaging data. Frontiers in Neuroscience 10. doi:10.3389/fnins.2016.00365

Rootes-Murdy, K., Gazula, H., Verner, E., Kelly, R., DeRamus, T., Plis, S., et al. (2022). Federated analysis of neuroimaging data: A review of the field. Neuroinformatics 20, 377–390. doi:10.1007/s12021-021-09550-7

Senanayake, N., Podschwadt, R., Takabi, D., Calhoun, V., and Plis, S. (2022). NeuroCrypt: Machine learning over encrypted distributed neuroimaging data. Neuroinformatics 20, 91–108. doi:10.1007/s12021-021-09525-8. Epub 2021 May 4

Thompson, P. M., Andreassen, O. A., Arias-Vasquez, A., Bearden, C. E., Boedhoe, P. S., Brouwer, R. M., et al. (2017). ENIGMA and the individual: Predicting factors that affect the brain in 35 countries worldwide. NeuroImage 145, 389–408. doi:10.1016/j.neuroimage.2015.11.057

Thompson, P. M., Stein, J. L., Medland, S. E., Hibar, D. P., Vasquez, A. A., Renteria, M. E., et al. (2014). The ENIGMA consortium: large-scale collaborative analyses of neuroimaging and genetic data. Brain Imaging and Behavior 8, 153–182. doi:10.1007/s11682-013-9269-5

Turner, J. A., Calhoun, V. D., Thompson, P. M., Jahanshad, N., Ching, C. R., Thomopoulos, S. I., et al. (2022). ENIGMA + COINSTAC: improving findability, accessibility, interoperability, and re-usability. Neuroinformatics 20, 261–275. doi:10.1007/s12021-021-09559-y

Vogt, N. (2023). Reproducibility in MRI. Nature Methods 20. doi:10.1038/s41592-022-01737-3

